# Predation by nematode-trapping fungus triggers mechanosensory-dependent quiescence in *Caenorhabditis elegans*

**DOI:** 10.1101/2024.05.14.594062

**Authors:** Tzu-Hsiang Lin, Han-Wen Chang, Rebecca J. Tay, Yen-Ping Hsueh

**Affiliations:** Genome and Systems Biology Degree Program, Academia Sinica and National Taiwan University, Taipei 106, Taiwan; Molecular Cell Biology, Taiwan International Graduate Program, Academia Sinica and Graduate Institute of Life Science, National Defense Medical Center, Taipei 114, Taiwan; Institute of Molecular Biology, Academia Sinica, Taipei 115, Taiwan

**Author notes:** Corresponding author Yen-Ping Hsueh Institute of Molecular Biology, Academia Sinica 128 Academia Road, Section 2, Nangang, Taipei 115, Taiwan.

**Keywords:** *C*. *elegans*/ mechanosensation/ nematode-trapping fungi/ predator-prey interactions/ sleep-promoting neuron

## Abstract

Predation can induce behavioral changes in prey, yet the molecular and neuronal mechanisms underlying prey responses remain poorly understood. Here, we investigated how the nematode *Caenorhabditis elegans* responds to predation by the nematode-trapping fungus, *Arthrobotrys oligospora*. We found that *A. oligospora* predation induced quiescence in *C. elegans* showing rapid cessation of pharyngeal pumping and movement. Calcium imaging revealed that this quiescence was regulated by the activation of sleep-promoting neurons, ALA and RIS. Genetic analyses demonstrated that ALA were essential for pharyngeal pumping inhibition, whereas both ALA and RIS contributed to movement cessation. Transcriptomic analysis in *C. elegans* showed the upregulation of immune defense genes in response to *A. oligospora* predation. We demonstrated that mechanosensation was required for pumping inhibition and transcriptomic regulation upon *A. oligospora* trapping. These findings suggest that physical constraints imposed by fungal traps trigger a stress-induced quiescence and the upregulation of defense genes in *C. elegans*. We suggest that trapping-induced quiescence might be a predation strategy used by sessile predators to prevail in the evolutionary arms race.

## Introduction

Organisms respond to stresses by altering their physiology and behavior. Abiotic stresses such as nutrient limitation, heat stress, and osmotic stress, tend to elicit highly conserved physiological responses in cells and organisms across the tree of life. For example, in *E. coli*, budding yeast, and human cells, when cells encounter heat stress, the heat shock transcription factor translocate into the nucleus to activate the transcription of heat shock proteins such as chaperones and other proteins that function to prevent protein misfolding and denaturation at high temperatures (Morimoto, 1998). In contrast, cellular responses to biotic stresses, arising from interactions such as pathogen infections, parasitism, competition, and predation, tend to be less conserved and more diverse (Baugh & Hu, 2020; Clarke *et al*, 2019; Hislop *et al*, 2007; Wilson *et al*, 2018). These responses are often specific to the organism’s evolution history and environmental context with a wide range of strategies against these challenges. Predation is one of the most pervasive and dramatic types of biotic stress, often resulting in a life-or-death scenario. Predation risk can lead to various responses in prey species, including a decline in foraging, reduced reproductive rates, and decreased motility (De Franceschi *et al*, 2016; Pribadi *et al*, 2023; Skelhorn, 2018; Stoks *et al*, 2003). Despite these diverse behavioral changes to predation risk, the molecular and neuronal mechanisms underlying these responses remain largely understudied.

The nematode *Caenorhabditis elegans* is an ideal model for studying the stress response at molecular and neuronal levels because it encounters diverse abiotic and biotic stresses such as temperature fluctuation, osmotic shock, starvation, infections, and predation in its natural environments (Cook *et al*, 2019; Frezal & Felix, 2015; Nigon & Félix, 2017). Notably, many of such environmental stresses induce a sleep-like behavior in *C. elegans* (Hill *et al*, 2014). It has been shown that stress-induced sleep in *C. elegans* is mediated through two peptidergic sleep-promoting interneurons, ALA and RIS (Hill *et al*., 2014; Iannacone *et al*, 2017; Nath *et al*, 2016; Nelson *et al*, 2014; Turek *et al*, 2013). Moreover, the epidermal growth factor receptor (EGFR) is expressed in ALA and RIS and is involved in stress-induce sleep regulation (Konietzka *et al*, 2020; Van Buskirk & Sternberg, 2007). Stress-induced sleep response is well-conserved among nematodes, flies, zebrafish, and mice, and this mechanism plays essential roles in increasing survival rates amidst different stress conditions (Kuo & Williams, 2014; Lee *et al*, 2019; Toda *et al*, 2019; Yu *et al*, 2022). Furthermore, a recent study has suggested that *C. elegans* antimicrobial peptides (AMPs) upregulated in response to pathogen infection and wounding, also regulate sleep (Sinner *et al*, 2021). In addition to their function in the pathogen defense response, AMPs assist in transmitting sleep signals from peripheral tissues to the nervous system, a mechanism observed in both *C. elegans* and *Drosophila* (Sinner *et al*., 2021; Toda *et al*., 2019).

Nematodes, including *C. elegans*, face predation stress from nematode-trapping fungi in their natural habitat. Fossil evidence of this predator-prey interaction between fungi and nematodes suggests a long evolutionary history, with amber fossils dating back 100 million years to the age of the dinosaurs (Schmidt *et al*, 2007). Our previous work has demonstrated that the interaction is still highly ecologically relevant today, as nematodes and nematode-trapping fungi are ubiquitous, and sympatric in over 60% of the randomly collected soil samples (Yang *et al*, 2020). In particular, *C. elegans* and the nematode-trapping fungus, *Arthrobotrys oligospora* had been isolated from the same sample collected in an apple orchard in Taiwan, indicating their coexistence (Yang *et al*., 2020). *A. oligospora*, despite being sessile, employs a range of adaptations to ensnare its nematode prey (de Ulzurrun & Hsueh, 2018); it attracts diverse species of nematodes by producing several volatile organic compounds (VOCs) that mimic nematode sex and food cues (Hsueh *et al*, 2017). When prey signals are present and nutrients are limited, *A. oligospora* forms three-dimensional adhesive networks to capture nematodes (de Ulzurrun & Hsueh, 2018). It has been shown that *A. oligospora* eavesdrops on nematode pheromones, ascarosides, via GPCRs (Hsueh *et al*, 2013; Kuo *et al*, 2024), which activate the conserved downstream signaling cascades such as the pheromone responsive MAPKs (Chen *et al*, 2020) and cAMP-PKA (Chen *et al*, 2022) pathways to trigger trap development. However, how *C. elegans* responds to predation by *A. oligospora* remains less clear.

In this study, we examined the behavior, neuronal, and physiological response of *C. elegans* upon *A. oligospora* predation and identified that the nematode-trapping fungus *A. oligospora* induces behavioral quiescence in captured *C. elegans*. This quiescent state is dependent on the activation of the sleep-promoting neurons. Furthermore, we demonstrate the involvement of mechanosensation in both behavioral quiescence and transcriptomic regulation triggered by *A. oligospora* predation. Taken together, our findings suggest that predation by *A. oligospora* induces mechanical stress on *C. elegans*, leading to quiescence and the upregulation of defense genes in this nematode.

## Results

### *A. oligospora* predation induces behavioral quiescence in *C. elegans*

To investigate how nematodes respond to predation stress caused by nematode-trapping fungi, we directly observed the behavior of *C. elegans* when the animals were trapped by *A. oligospora*. Notably, *C. elegans* exhibited an immediate intense struggling movement, but could not escape from the three-dimensional adhesive traps of the fungus. Interestingly, *A. oligospora*-trapped animals ceased their movement after 15–20 minutes (Movie EV1). To better define the nature of this behavior, we measured and quantified the three distinct features of *C. elegans* quiescence: pharyngeal pumping, movement, and response to stimuli in *A. oligospora*-trapped *C. elegans*. We found that the pharyngeal pumping was inhibited by *A. oligospora*-trapping, with the pumps per second dropping dramatically from 4.8 to 1.3 Hz within 5 minutes and completely stopped at approximately 20 minutes (Fig. 1A). We next measured the head struggle movement in trapped animals and found that most animals continuously struggled for ∼20 minutes and then entered a quiescent state (Movie EV1 and Fig. 1B). We next measured the animal’s response to stimuli by applying glycerol as the aversive stimulus to the head of the animals and monitored its avoidance response (Hilliard *et al*, 2002). We found that both free-moving and *A. oligospora*-trapped animals that still exhibited the struggling behavior showed a strong avoidance response to glycerol (Fig. 1C). In contrast, *C. elegans* that had entered a quiescent state after being trapped for 15 minutes or longer (30 minutes) displayed a strong reduction in the avoidance to glycerol (Fig. 1C). Taken together, our results demonstrate that *A. oligospora* predation triggers the cessation of pharyngeal pumping and movement as well as a decreased ability to respond to external stimuli, representing three key features of the quiescence state in *C. elegans*.

**Figure 1.**
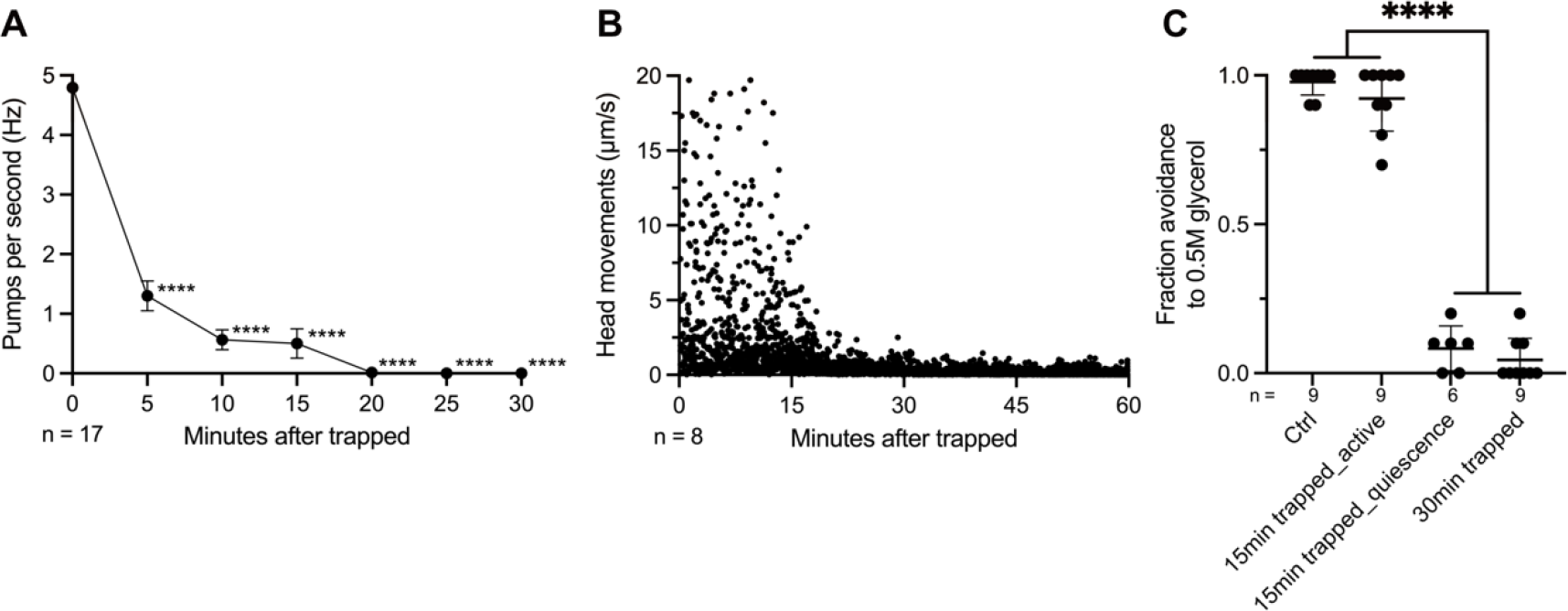
*A. oligospora* predation-induced quiescence in *C. elegans* (A) Pharyngeal pumping rates of WT after *A. olisospora* trapping. (B) Head movements of WT after *A. olisospora* trapping. (C) Response to aversive 0.5M glycerol of WT at different time points and behavior states after *A. olisospora* trapping. Mean ± SEM; One-way ANOVA, Tukey’s multiple comparison test; ****p < 0.0001.

### The sleep-promoting neurons ALA and RIS are activated in response to *A. oligospora* trapping

In *C. elegans*, two sleep-promoting neurons, ALA and RIS, are crucial players in the sleep circuit (Turek *et al*., 2013; Van Buskirk & Sternberg, 2007). To determine whether *A. oligospora* trapping in *C. elegans* also activates these two neurons, we imaged the neuronal activity of ALA and RIS neurons in trapped nematodes by monitoring calcium levels using GCaMP. Our data revealed that neuronal activity reached maximal levels, with a 72% increase in ALA neurons and a 169% surge in RIS neurons approximately 20 minutes after being trapped (Fig. 2A,C). This timing coincided perfectly with the behavioral quiescence observed (Fig. 1), suggesting that the activation of ALA and RIS upon trapping triggers the quiescence response of *C. elegans*.

**Figure 2.**
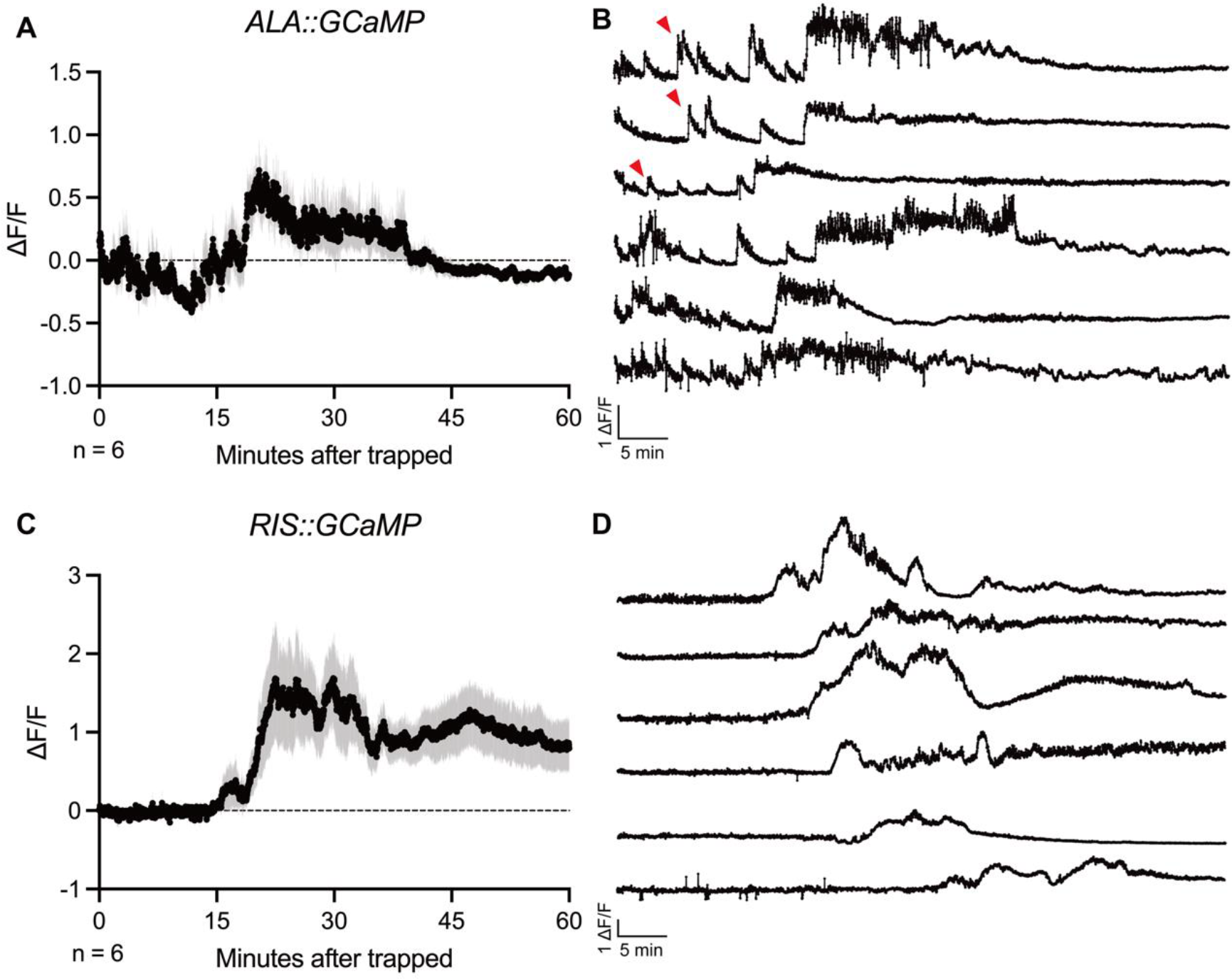
Sleep-promoting neurons ALA and RIS are activated in 20 min after trapping (A) Average trace of the ALA calcium response to *A. oligospora* trapping. Mean ± SEM (B) Individual traces of the ALA calcium response to *A. oligospora* trapping. Arrows indicate the early peak response in the first 20 min. (C) Average trace of RIS calcium response to *A. oligospora* trapping. Mean ± SEM (D) Individual traces of the RIS calcium response to *A. oligospora* trapping.

Notably, ALA and RIS neurons exhibited distinct firing patterns in animals trapped by *A. oligospora*. The ALA neurons exhibited a dense (at least 3 peaks) but mild (41% of the maximal activation level at ∼20 min) firing soon after being trapped by *A. oligospora* (Fig. 2B), coinciding with the observed rapid pumping inhibition. ALA then reached its maximal activation around 20 min after trapping, and the calcium level gradually returned to baseline after another 20 minutes (Fig. 2A). In contrast, RIS neurons showed steady baseline calcium levels for the first 20 minutes after trapping and depolarization to the maximal value, coinciding with the timing of movement cessation (Fig. 2D). Furthermore, the neuronal activity persists for an ensuing 40 minutes until the end of the imaging session (Fig. 2C). These results suggest that both sleep-promoting neurons are maximally activated at 20 min after trapping, coinciding with the movement quiescence behavior observed in trapped animals. The distinct firing patterns of ALA and RIS could potentially contribute to the temporal regulation of pumping inhibition and movement cessation observed in *A. oligospora*-trapped *C. elegans*.

### The sleep-promoting neurons regulate pharyngeal pumping and movement inhibitions

Studies have shown that CEH-14, a LIM class homeodomain protein, is required for ALA-specific gene expression and axon outgrowth (Van Buskirk & Sternberg, 2010), whereas APTF-1, an AP2 transcription factor, is required for RIS function (Turek *et al*., 2013). To investigate the functional roles of ALA and RIS neurons in pumping inhibition and movement cessation upon *A. oligospora* trapping, we compared the behavior of *ceh-14(ch3)* (ALA-deficient) and *aptf-1(gk794)* (RIS-deficient) mutants to that of the wild-type. In wild-type animals, a dramatic decrease in pharyngeal pumping rate, plummeting from 4.7 Hz to 0.6 Hz was observed in the first initial 5 minutes after trapping (Fig. 3A), whereas in *ceh-14*, there was only a slight reduction in pumping rate from 4.7 Hz to 3.8 Hz during the same period (Fig. 3A). Across the entire time assayed, *ceh-14* showed a more gradual decrease in pumping rate compared to the WT (Fig. 3A). In contrast, the *aptf-1* mutants showed the same degree of pumping inhibition as WT animals, and the *ceh-14 aptf-1* double mutant did not intensify the suppression of pumping inhibition beyond that observed in the *ceh-14* mutant alone (Fig. 3A). These results, together with the observed calcium activation pattern within the initial 20 minutes of trapping (Fig. 2B), suggest that ALA neurons encode the neuronal signals that inhibit pharyngeal pumping upon *A. oligospora* predation, and that RIS is not required for trapping-induced pharyngeal pumping cessation (Fig. 3A).

**Figure 3.**
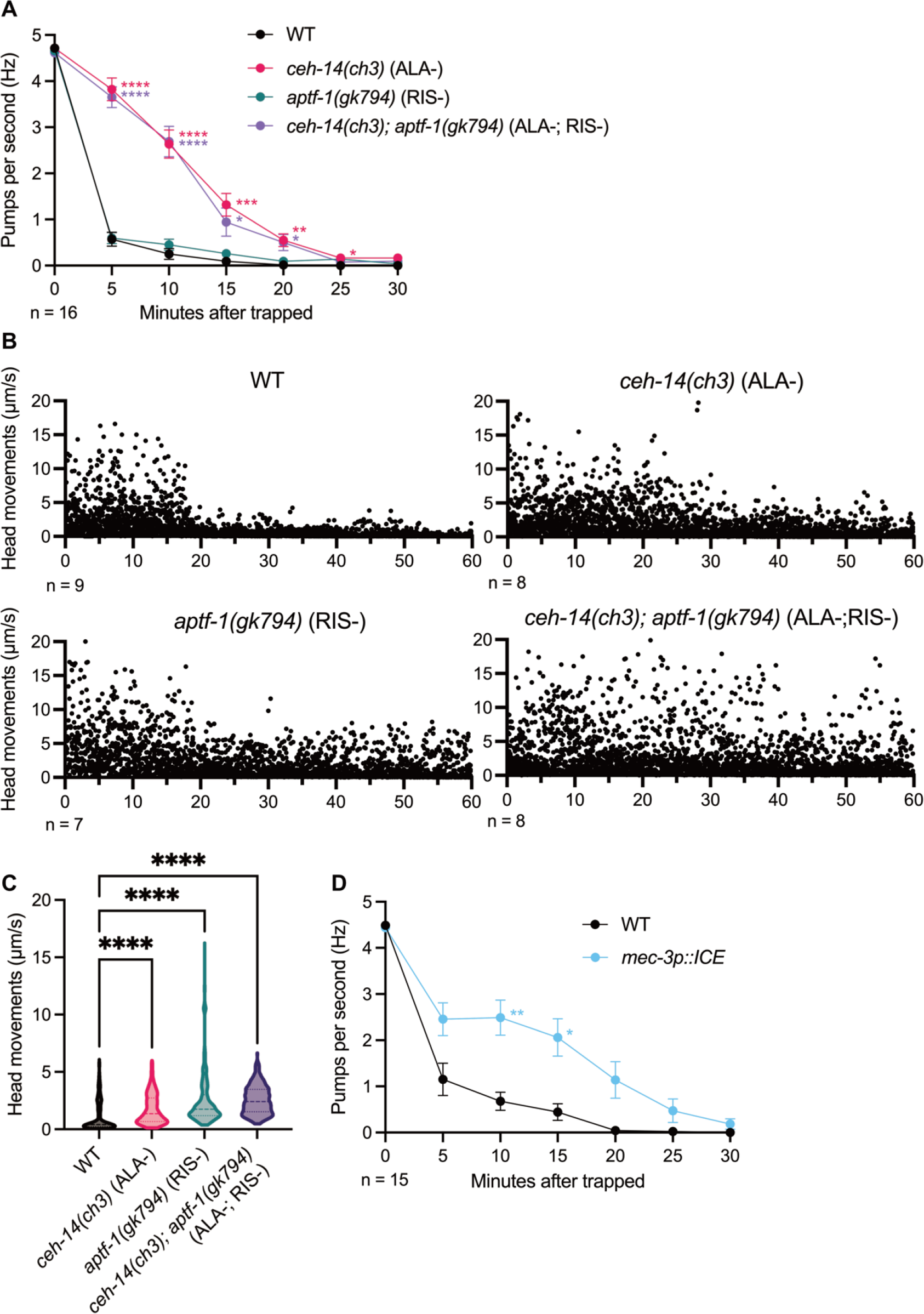
Trapping-induced quiescence is regulated by sleep-promoting neurons and mechanosensory neurons (A) Pharyngeal pumping rates of wild-type, ALA-deficient, RIS-deficient, and ALA/RIS- deficient mutants after *A. olisospora* trapping. (B) Head movements of wild-type, ALA-deficient, RIS-deficient, and ALA/RIS-deficient mutants after *A. olisospora* trapping. (C) Distribution of head movements of wild-type, ALA-deficient, RIS-deficient, and ALA/RIS- deficient mutants in Figure 3B. (D) Pharyngeal pumping rates of genetically ablated mechanosensory neurons after *A. olisospora* trapping. (A and D) Mean ± SEM; Two-way ANOVA, Šidák’s multiple comparison test, comparing strain effects at each timepoint; *p < 0.05, **p < 0.01, ***p < 0.001, ****p < 0.0001. (C) Kruskal-Wallis test with Dunn’s correction; ****p < 0.0001.

Next, we investigated whether these sleep-promoting neurons played a role in the cessation of movement induced by *A. oligospora* predation. We measured the head movement of wild-type, *ceh-14*, *aptf-1,* and *ceh-14; aptf-1* animals upon *A. oligospora* trapping. Contrary to what we observed in pumping inhibition, we found that both ALA and RIS deficient mutants were more active than the wild-type after being trapped by *A. oligospora* (Fig. 3B,C). These data suggest that both ALA and RIS neurons both regulate the movement cessation triggered by *A. oligospora* trapping.

### Mechanosensory neurons regulate trapping-induced pharyngeal pumping

Next, we asked which signals from *A. oligospora* traps triggered the sleep response. A previous study reported that physical restriction in a microfluidic device can trigger the sleep-like state of *C. elegans* through the mechanosensory pathway and RIS activation (Gonzales *et al*, 2019). Therefore, we hypothesized that trap adhesion immobilizing nematodes might represent a mechanical stimulus to trigger quiescence. To test this hypothesis, we mimicked trap adhesion by physically constraining the animals with WormGlu. Nematodes were glued with brief cold immobilization and recovered at room temperature. Pharyngeal pumping was completely abolished in response to the cold shock. After temperature-shifted recovery, the glued *C. elegans* failed to restore pharyngeal pumping, while the control animal resumed pumping in 5 minutes, indicating that physically constraining the animals with WormGlu was sufficient to trigger the cessation of pharyngeal pumping (Fig. EV1A). To further define the role of the mechanosensory pathway in *A. oligospora*-induced quiescence, we next generated a mutant in which the mechanosensory neurons (ALML/R, AVM, FLP, PLML/R, PVD, and PVM) were genetically ablated by the expression of human caspase ICE under the promoter *mec-3p::ICE*. We then measured the pharyngeal pumping of the mechanosensory-ablated mutant upon *A. oligospora* trapping and found that the mutant exhibited a more moderate reduction in pharyngeal pumping than the wild-type (Fig. 3D). These results suggest that the mechanosensory pathway contributes, at least partially, to trapping-induced quiescence.

Next, we investigated whether mechanosensory pathways play a role in trapping-induced sleep circuit activation. To this end, we used calcium imaging to measure the activity of ALA and RIS upon *A. oligospora* predation in the mechanosensory-deficient mutants. In mutants in which 10 of the 30 mechanosensory neurons were genetically ablated, the early ALA activity patterns in the first 15 min were different from those in wild-type animals. Specifically, the short and dense peaks observed in the wild-type ALA tracks (Fig. 2B) were reduced in the mechanosensory-ablated mutants (Fig. EV2A,B). In contrast, the activation of RIS neurons was only slightly reduced, although this was not statistically significant (Fig. EV2C,D). The partial reduction in the activity of ALA and RIS neurons suggests that additional mechanosensory neurons might still participate in trapping-triggered quiescence or that additional chemical cues from *A. oligospora* could trigger *C. elegans* quiescence in synergy with mechanical stress.

### *A. oligospora* trapping induces up-regulation of defense and immune response genes

To identify the transcriptomic changes in *C. elegans* in response to *A. oligospora* predation, we conducted RNA sequencing on nematodes that were trapped by *A. oligospora* and those freely moving on a fungal lawn for 30 and 60 min (Fig. 4A). We found 350 and 613 upregulated differentially expressed genes (DEGs), and 24 and 129 downregulated DEGs in the 30- and 60-minute trapping RNA-seq, respectively (fold change > 2; p < 0.05) (Fig. 4B,C and EV3A). Gene Ontology enrichment analysis of the DEG datasets underscored that in the 30-minute condition, the top 14 functional categories for upregulated genes were all associated with defense and immune responses (Fig. 4D). Similar results were also observed in the 60-minute upregulated DEG datasets (Fig. EV3B). We observed similar upregulation patterns in defense and immune response-related genes across both the 30- and 60-minute trapped groups. (Fig. 4D). Notably, p38 MAPK pathway genes like *tir-1*, *dlk-1*, *pmk-3*, *cebp-1*, and *vhp-1*, were upregulated and known to regulate innate immunity and axon regeneration (Fig. 4B and EV3A). Additionally, a significant upregulation (2-fold to 38-fold) was noted in antimicrobial peptides (AMPs), including *cnc-2, 4, 10, 11*, and *nlp-8, 25, 27-34, 62*, in the 60-minute trapped group (Fig. 4B and EV3A).

**Figure 4.**
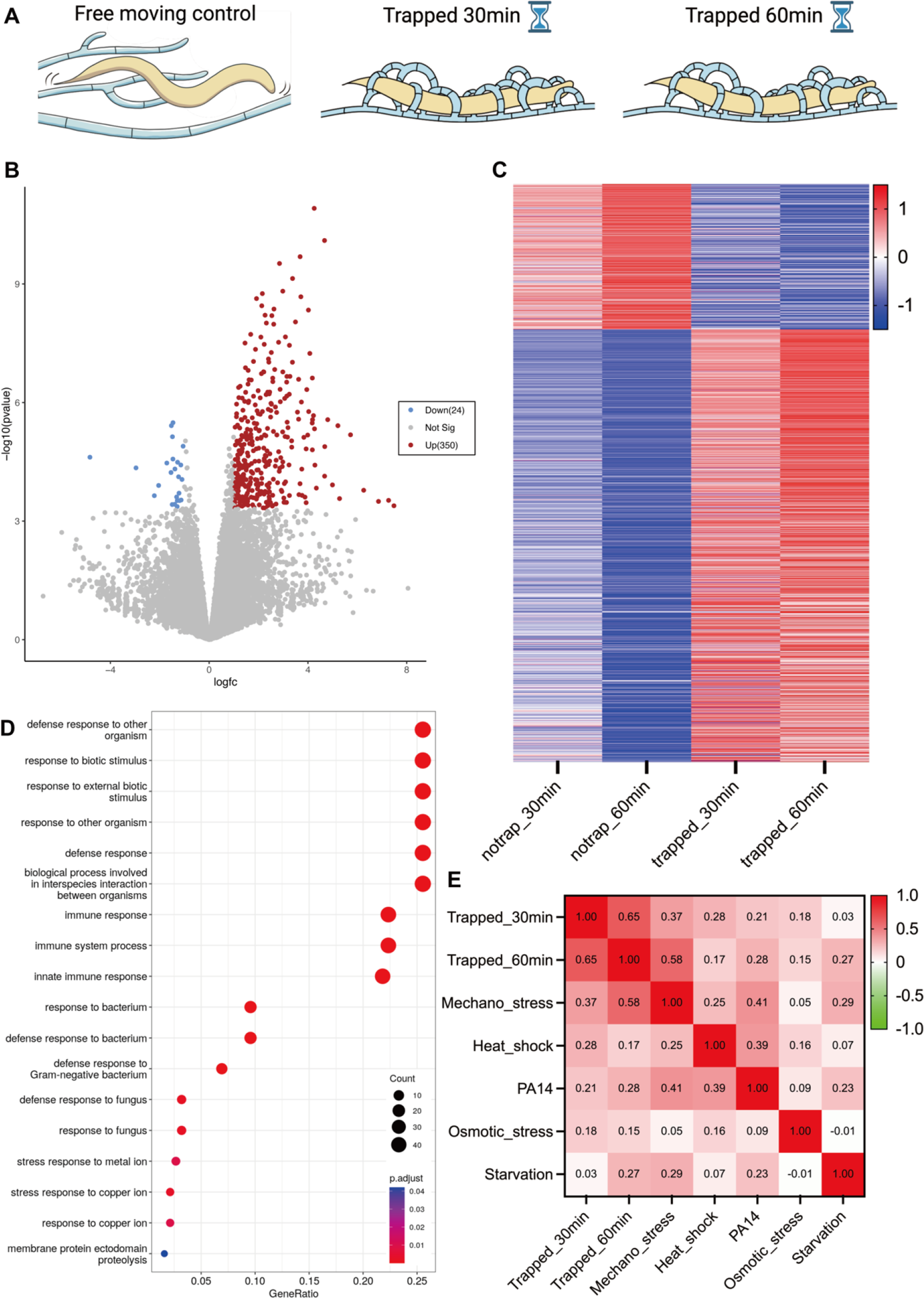
Up-regulation of immunity genes and the high correlation between trapping- and mechano-induced transcriptomic change (A) Schematic of the experimental conditions used for RNA-seq. (B) Volcano plot of RNA-seq analysis of 30 min trapped groups compared with the no-trap control. Threshold fold-change > 2; p-value < 0.05. (C) Heat map showing the expression patterns of differentially expressed genes in the different experimental groups. Normalized in Z-scale. (D) Plots of the gene enrichment analysis from GO molecular functions of 30 min trapped groups compared with the no-trap control. (E) Heat map of the cross-correlation matrix with Spearman’s ρ comparing different published stress-induced transcriptomic data.

Pathogens can release toxins that trigger host cells to activate detoxification pathways. These pathways involve enzymes, such as Cytochrome P450 (CYP450), UDP-glucuronosyltransferase (UGT), glutathione S-transferase (GST), and ATP-binding cassette (ABC) transporters, which work together to metabolize and eliminate xenobiotics. Although a minimal number of *cyp*-family genes were downregulated in the trapped conditions, the majority of *cyp*, *ugt*, *gst*, and ABC transporters were upregulated in both the 30- and 60-minute trapped groups (Fig. EV3C). We found that the transthyretin-related family (TTR), a nematode-specific expanded protein family, was overrepresented among the upregulated genes (Fig. EV3C). One of the most significantly upregulated TTR genes, *ttr-33*, has been reported to have a protective role against oxidative stress, as well as a neuroprotective role (Offenburger *et al*, 2018). Our data suggest that *A. oligospora* trapping induced a majority of upregulated genes, which comprise immune regulating and defense response genes.

### Trapping-induced transcriptomic regulations are highly correlated with mechanical stress

We have demonstrated that mechanosensory neurons play a role in *A. oligospora* trapping-induced sleep. To determine whether mechano-signal perception influences transcriptional changes, we compared our RNA-seq data from *A. oligospora* trapping with published transcriptomic datasets from *C. elegans* exposed to diverse stresses. These stresses include heat shock (McColl *et al*, 2010), *Pseudomonas aeruginosa* (PA14) infection (Miller *et al*, 2015), osmotic stress (Rohlfing *et al*, 2010), starvation (Uno *et al*, 2013), and mechanical trauma (Egge *et al*, 2021), all of which *C. elegans* may encounter in its natural environment. Cross-correlation analysis was performed on the transcriptomic data using Spearman’s rank correlation coefficient. Our cross-correlation analysis revealed a high correlation (ρ=0.65) between the transcriptomic profiles of *C. elegans* trapped for 30 minutes and 60 minutes, which was expected (Fig. 4E). Surprisingly, the Spearman’s ρ between the mechanical trauma datasets and *A. oligospora*-trapped 30-minute datasets was 0.37, and this correlation coefficient increased to 0.58 comparing to the *A. oligospora*-trapped 60-minute datasets (Fig. 4E). Additionally, we compared the upregulated DEGs between mechanically induced (Egge *et al*., 2021), trapping-induced 30 min, and trapping-induced 60 min RNA-seq. We found that 182 core genes were consistently upregulated across all three RNA-seq datasets comprising antimicrobial peptides of the *cnc* and *nlp* families, both members of *igeg* (Immunoglobulin, EGF, and transmembrane domain), and genes that respond to biotic stimulus (GO:0009607), biological process involved in interspecies interaction between organisms (GO:0044419), and defense response (GO:0006952) (Table EV1). Furthermore, 198/350 (57%) and 335/613 (55%) upregulated DEGs in the trapping-induced 30 and 60 min, respectively, were shared with upregulated genes from the mechanically induced dataset (Fig. EV3D). These results suggest that *A. oligospora* trapping and mechanical stress trigger overlapping pathways that regulate the transcriptomic landscape in *C. elegans*.

### Reactive oxygen species and epidermal growth factor receptor signaling pathways for trapping-induced sleep

Next, we investigated the signaling factors responsible for trapping-induced sleep. Studies have suggested that AMPs can play a dual role in both innate immunity and sleep promotion (Sinner *et al*., 2021; Toda *et al*., 2019). Given this, we tested the impact of AMPs on the *A. oligospora* trapping-induced sleep response in the strain PHX1446, in which 19 AMP members of the *nlp* and *cnc* families were knocked out. However, we did not observe any significant effect on pumping inhibition in PHX1446 worms (Fig. EV4A).

Previous studies have indicated that reactive oxygen species (ROS) can inhibit pharyngeal pumping in response to UV irradiation (Bhatla & Horvitz, 2015). In addition, our finding that xenobiotic metabolism genes, which are commonly associated with ROS, are upregulated in response to trapping and the predicted role of the *ttr* gene family in oxidative stress led us to hypothesize that ROS may act as a signaling factor for *A. oligospora*-induced sleep. To test this hypothesis, we imaged the real-time H_2_O_2_ levels in worms trapped by *A. oligospora* using the genetically encoded H_2_O_2_ sensor HyPer under the ribosomal promoter *rpl-17* (Fig. 5A). We observed that the concentration of H_2_O_2_ gradually increased over 15 min to its peak level, before decreasing slightly thereafter (Fig. 5A). It is interesting to note that ROS levels reach their maximum immediately before nematodes enter their quiescent state. To confirm that ROS are involved in signal transduction, we applied the well-established antioxidant N-acetylcysteine (NAC) to *C. elegans* before *A. oligospora* trapping. Our results showed prolonged pharyngeal pumping, indicating that ROS contributes to *A. oligospora*-trapping induced quiescence behavior (Fig. 5B).

**Figure 5.**
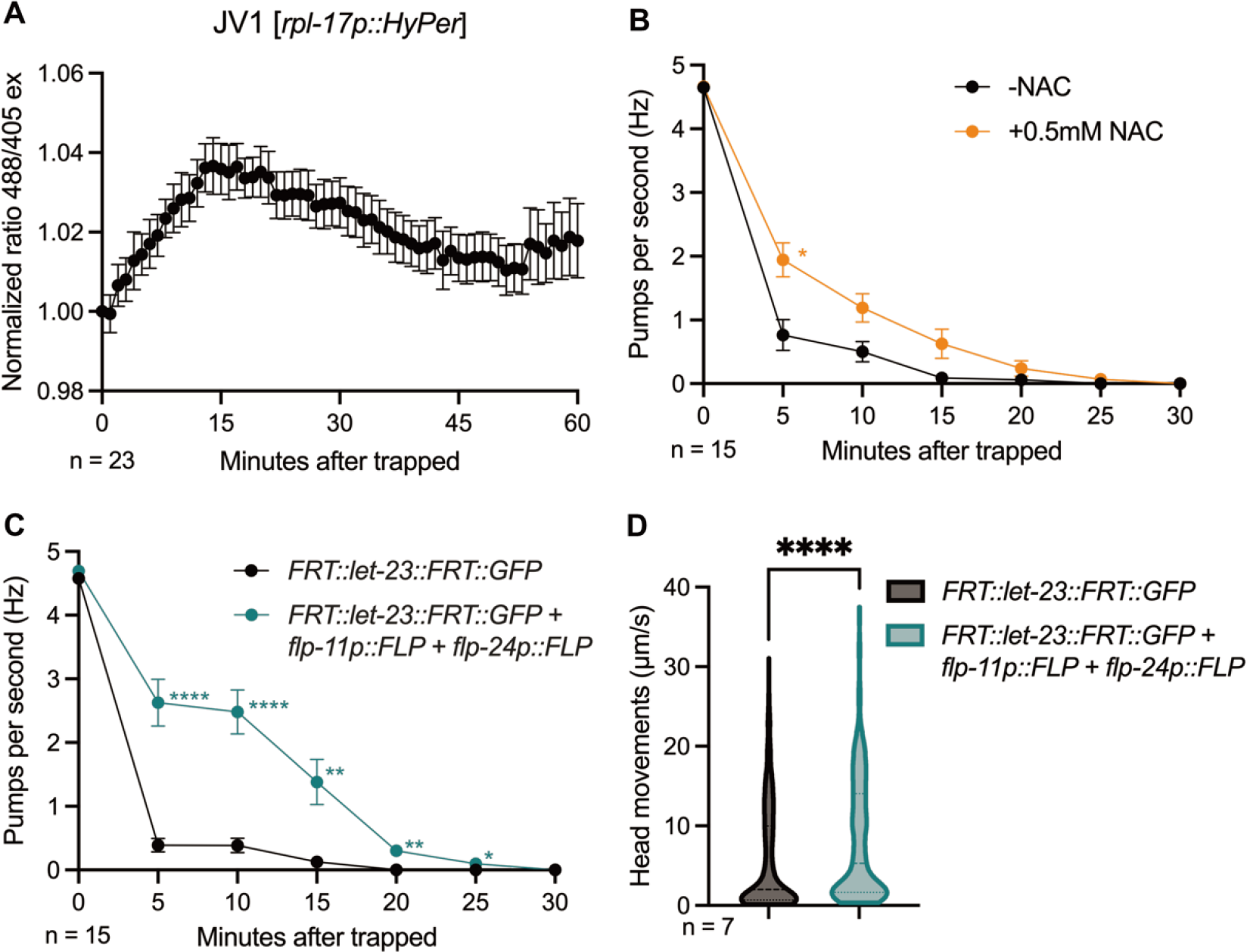
ROS and EGFR signaling pathways are involved in trapping-induced quiescence (A) A Normalized ratio at 488/405 excitation indicates oxidative levels sensed by HyPer in response to *A. oligospora* trapping. Mean ± SEM. (B) The effects of antioxidant NAC treatment on trapping-induced pumping inhibition. (C) Pharyngeal pumping rates of *let-23* knockdown specifically in ALA and RIS after *A. olisospora* trapping. (D) Distribution of head movements in the WT and *let-23* knockdown mutants. (B and C) Mean ± SEM; Two-way ANOVA, Šidák’s multiple comparison test, comparing strain effects at each timepoint; *p < 0.05, **p < 0.01, ***p < 0.001, ****p < 0.0001. (D) Mann–Whitney test; ****p < 0.0001.

Stress-induced sleep in *C. elegans* involves the epidermal growth factor receptor (EGFR) signaling pathway, the *C. elegans* homolog of EFGR LET-23 is expressed in both sleep-promoting neuron ALA and RIS (Konietzka *et al*., 2020). It had been shown that one can employ conditional knockdown of *let-23* by FLPase-induced recombination to study the role of *let-23* in heat shock (Konietzka *et al*., 2020). Here, we used ALA and RIS specific *flp-24* and *flp-11* promoters to drive FLP expression to knockdown *let-23*. We found that disrupting the EGFR pathway, specifically in sleep-promoting neurons, ALA and RIS, led to prolonged pharyngeal pumping and movement during *A. oligospora*-induced quiescence (Fig. 5C,D), indicating that the EGFR pathway plays an important role in *A. oligospora* predation-triggered quiescence.

## Discussion

Predators play a critical role in shaping the behavior of prey at both immediate and evolutionary time scales. In the realm of terrestrial predator-prey dynamics, most interactions are typified by a “catch and run” scenario, emphasizing motility as a core strategy for both predator and prey. For example, physical restraint by predators often triggers death feigning in insects, fish, birds, and mammals, such behavior might trick the predators to loosen their bite (Humphreys & Ruxton, 2018; Skelhorn, 2018). Damselfly larvae and sticklebacks decrease mobility and reproduction in the presence of predators by hiding strategy (Candolin, 1998; Stoks *et al*., 2003). Mice elicit freezing behavior to avoid detection from avian predators (De Franceschi *et al*., 2016). These anti-predator behaviors of transition from motile to quiescent allow prey animals to avoid detection by active predators.

On the other hand, sessile predators have evolved multiple alternative predation strategies that circumvent the need for a chase. A notable example of such adaptation can be found in carnivorous plants, such as the Venus flytrap, which releases volatile organic compounds (VOCs) including terpenes, to lure naive prey (Kreuzwieser *et al*, 2014). Similarly, nematode-trapping fungus *A. oligospora* also secretes VOCs that mimic sex and food smell to lure their prey nematodes (Hsueh *et al*., 2017). We speculate that the induction of sleeping behavior in nematodes by *A. oligospora* trapping could serve as an additional strategy to enhance the likelihood of successful predation. Trap-induced sleep reduces the possibility of nematodes escaping from the predatory fungi, and thus, benefits the predators. As a result, predation-induced sleep might be a strategy that benefits sessile predators by avoiding prey evasion.

Stress-induced sleep is a well-defined response observed across diverse species (Kuo & Williams, 2014; Lee *et al*., 2019; Toda *et al*., 2019; Yu *et al*., 2022). In *C. elegans*, the sleep-promoting neurons mediated by the EGFR pathway trigger quiescent states when the organism encounters bacterial toxins, heat shock, or wounding (Hill *et al*., 2014; Konietzka *et al*., 2020; Sinner *et al*., 2021). Our study demonstrates that nematode-trapping fungus also triggers quiescence in *C. elegans* by engaging the same sleep-promoting neural circuit, suggesting that the predatory fungi co-opted this stress-induced sleep response for its own benefit. Sleep-promoting neurons, ALA and RIS, play a central role in predator-induced quiescence, exhibiting distinct temporal dynamics. ALA depolarization occurred within 5 minutes, while RIS depolarization reached a maximum at 20 minutes. This pattern aligns with previous findings on heat shock-induced quiescence, which suggested a potential hierarchy with ALA acting upstream of RIS (Konietzka *et al*., 2020). Our study revealed that disrupting sleep circuits does not completely abolish quiescence. This suggests that while these circuits are crucial for initiating behavioral quiescence, additional mechanisms likely also contribute to nematode quiescence induced by fungal predation. We hypothesized that after capturing *C. elegans* via adhesive nets, *A. oligospora* might secrete effectors that could help to maintain the nematodes in a quiescent and vulnerable state. Time-course transcriptional profiling of *A. oligospora* shows highly upregulated secretion-related genes after trapping nematodes (Lin *et al*, 2023). Recent works in *Arthrobotrys flagrans*, another fungal predator, also demonstrated that a fungal effector, CyrA, played a role in the full virulence of the fungus (Wernet *et al*, 2021). In parallel to our findings with sleep circuits, *A. flagrans* induce upregulation of *C. elegans nlp-27* to trigger neurodegeneration and paralysis (Pop *et al*, 2024). NLP-27 appears to contribute but may not be solely responsible for complete paralysis during fungal infection (Pop *et al*., 2024). This support the hypothesis that multiple mechanisms are integrated to regulate nematode response during fungal predation.

Mechanosensory neurons make up 10% of *C. elegans* nervous system, indicating the essential role of mechanosensation in this organism (Goodman, 2006; Goodman & Sengupta, 2019). A previous study on a different nematode-trapping fungus, *Drechslerella doedycoides*, which catches nematodes with its constricting hyphal rings, demonstrated that when a nematode touches a trap, it triggers a rapid reversal response withdrawing from fungal traps (Maguire *et al*, 2011). Touch-insensitive *mec-4* mutants lead to a greater capture rate, emphasizing the importance of a functional mechanosensory circuit crucial for escaping from fungal predators (Maguire *et al*., 2011). Here, we show that mechanosensation also plays a key role in trapping-induced quiescence. Moreover, the transcriptional response of nematode-trapping fungus predation recapitulated the upregulation of defense immune genes in *C. elegans* under mechanical stress. These lines of evidence demonstrate that mechanosensory circuits have a multifaceted role in different aspects of predator responses.

Taken together, our results reveal that predation by nematode-trapping fungus induces mechanosensory-dependent sleep and transcriptomic changes in *C. elegans*. Activation of sleep-promoting neurons regulates pharyngeal pumping, movement inhibition, and upregulation of defense immune genes in nematode prey (Fig. 6). This work sheds light on the neuronal and genetic mechanisms underlying the *C. elegans* response to fungal predation and provides insight into the understanding of predation-triggered behavioral changes in an animal.

**Figure 6.**
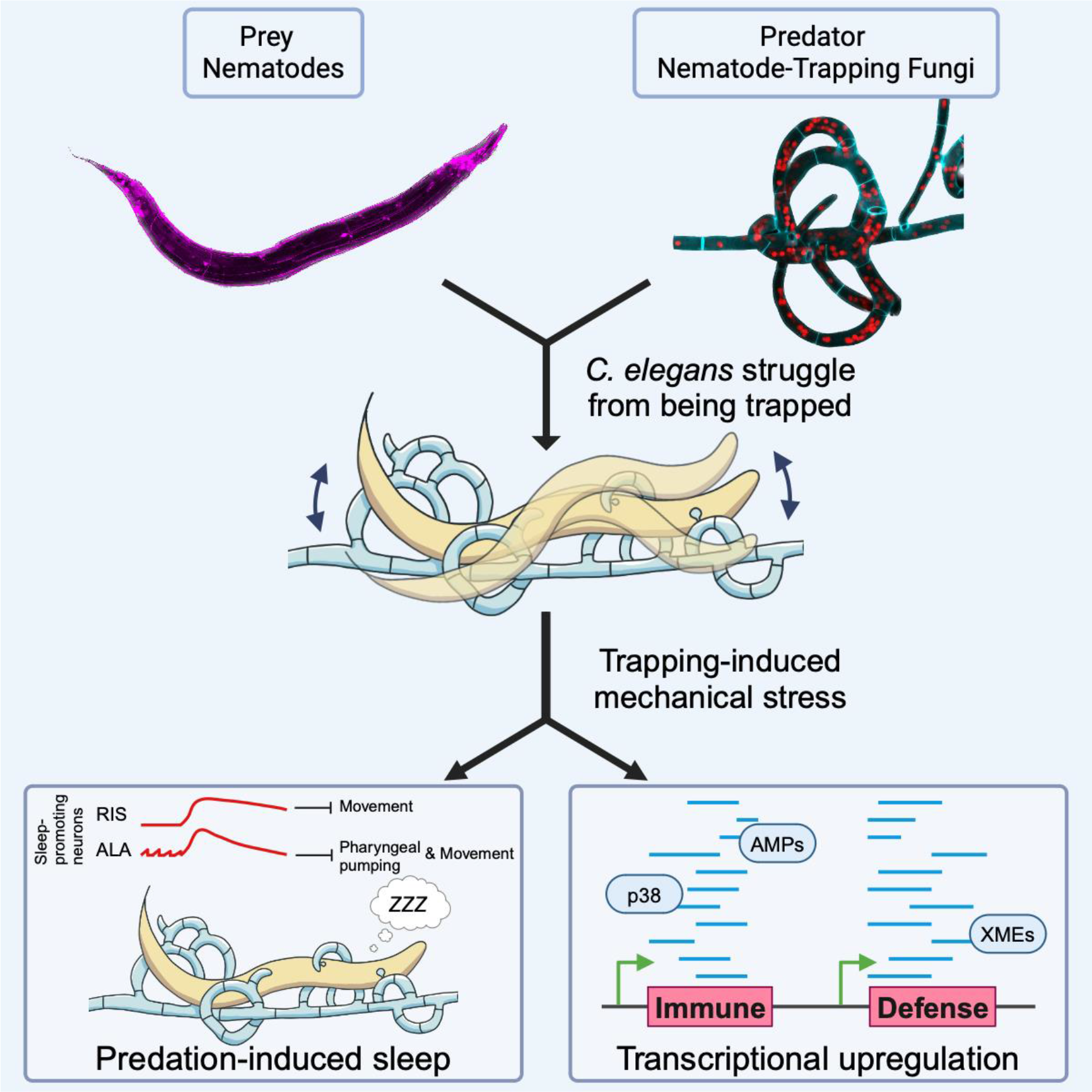
A model for *A. oligospora*-trapping induced sleep in *C. elegans*

## Methods

### Methods and protocols

#### Strains and culture conditions

The wild-type strain was the *C. elegans* strain CGC1. *C. elegans* were grown at 23°C on nematode growth media (NGM) plates seeded with bacteria (*E. coli* OP50) as a food source. The sex and age of the animals used for each experiment were adult hermaphrodites. All transgenic strains used in this study are listed in the SI Appendix, Table S1. *Arthrobotrys oligospora* strain TWF154 was used in this study. For culture maintenance, *A. oligospora* was grown at 25°C on potato dextrose agar (PDA, i.e., rich-nutrient) plates. All experiments were conducted on low-nutrient medium (LNM:2% agar, 1.66 mM MgSO_4_, 5.4 µM ZnSO_4_, 2.6 µM MnSO_4_, 18.5 µM FeCl_3_, 13.4 mM KCl, 0.34 µM biotin, and 0.75 µM thiamin) at 23°C.

#### Arthrobotrys oligospora trap induction

*Arthrobotrys oligospora* strain TWF154 was routinely maintained on PDA plates. Trap induction was required before conducting the experiments. A 3 mm square PDA fungal culture was chunked onto low-nutrient medium (LNM) for growth at 25°C for 5 days as the pre-starved culture. A 3 mm square LNM chunk (pre-starved culture) was chunked onto LNM for growth at 25°C for 4 days. On day 4, when fungal hyphae had grown to the edge of the 5 cm plates, approximately 300-500 young adults/adults WT *C. elegans* were added to induce traps. Worms were incubated with *A. oligospora* overnight at 25°C to induce traps. All experiments were conducted on day 5 trap-induced cultures.

#### Quantifying pharyngeal pumping

Adult hermaphrodite *C. elegans* were washed with M9 buffer and gravity-settled for 1-2 min to avoid pick and centrifuge stress. One hundred worms were transferred to trap-induced *A. oligospora* plates and allowed to be trapped; typically, the nematodes would be trapped within a few minutes. Pharyngeal pumping was scored by the eye using a stereo-dissecting microscope for 15 sec every 5 min. Pharyngeal pumping rates were calculated by transforming the pumping number in 15 sec to Hz.

#### Tracking head movements

Adult hermaphrodite *C. elegans* were prepared with M9 buffer and gravity-settled as described in the previous section. We used *C. elegans* expressing a fluorescent marker in the pharyngeal muscle to create high-contrast images for automatic tracking. Time-lapse videos were captured using a stereo-dissecting microscope at 6 frames per minute for 1 hour with continuous GFP excitation. Videos were analyzed by Tracker (Open Source Physics) using an autotracker function with default settings and manually curated when objects were not detected.

#### Head-drop avoidance assay

We performed the avoidance assay using 0.5M glycerol on both active worms and trapping-induced quiescent worms. The nematodes to be tested were transferred by picking from LNM plates to non-seeded NGM plates. The drop assay for active worms was modified from the standard method described by Hilliard et al.(Hilliard *et al*., 2002) Briefly, 0.5M glycerol was dropped by a capillary tube in front of forward-moving worms and then sourced the number of positive backward movement when contacting the glycerol drop. For quiescent worms, a 0.5M glycerol drop was placed just ahead of the worms. Fraction avoidance was calculated as follows: (number of positive backward nematodes) / (total number of nematodes tested). Each dot represents the fraction avoidance of 10 tested worms.

#### Neuronal calcium imaging

GCaMP was used to monitor neuronal calcium levels in *C. elegans* ALA (*let-23p*) and RIS (*flp-11p*) neurons. After the worms were trapped by *A. oligospora*, a 3 cm LNM square with a tested worm at the center was chunked, inverted, and mounted on a cover glass. Imaging was performed using a Zeiss Axio Observer.D1 microscope with Photometrics Evolve 512 EMCCD camera at 1 fps for 60 minutes. Analyses were performed using Fiji to measure the fluorescence intensity of GCaMP in the region of interest (ROI). The fluorescence intensity in the first 1 min of each video was averaged to obtain the baseline. Data were calculated and normalized as (fluorescence intensity at each frame – baseline) / (baseline).

#### WormGlu physical restriction

To apply WormGlu to nematodes, we cold-immobilized worms by placing the plates on ice for 3 min. After the worms stopped moving, WormGlu was continuously applied to the worm surface for 2 minutes using a hand-pulled capillary. The assay was completely done on ice. *C. elegans* were recovered at 23°C for 5 min before scoring pharyngeal pumping for 30 min.

#### RNA-seq transcriptomic analysis

To investigate the response of *C. elegans* to *A. oligospora* predation, RNA-seq was used to identify the genes affected after 30 and 60 min of trapping. We harvested the nematodes that were either trapped (test groups) or allowed to crawl freely on a fungal lawn (control groups) for 30 and 60 min, respectively. Further analyses, including DEG and GO-term enrichment analyses, were based on RNA-seq experiments. Total RNA was extracted using the Trizol-chloroform method and treated with DNase followed by ethanol precipitation. RNA libraries for RNA-seq were prepared using Illumina TruSeq® Stranded mRNA Library Prep following the manufacturer’s protocol. RNA-seq analysis was performed using the STAR-RSEM-edgeR pipeline. Sequence reads were mapped to c_elegans.PRJNA13758.WS273 using Spliced Transcripts Alignment to a Reference (STAR). Read quantification was performed using RNA-Seq by Expectation-Maximization (RSEM). Gene expression normalization to logCPM was performed using Empirical Analysis of Digital Gene Expression Data in R (edgeR).

#### Cross-correlation analysis

Cross-correlation analysis was used to identify the relationship between the different stress-induced transcriptomic data. We used the 30-minute trapped DEGs as the genes of interest and calculated the fold-change of these genes from published stress-induced datasets, including heat shock (GSE22383), PA14 infection (GSE72029), osmotic stress (GSE19310), starvation (GSE27677), and mechanical trauma (GSE148325). Spearman’s correlation matrix was calculated using R version 4.3.0.

#### CRISPR-Cas9 knockout

The CRISPR STOP-IN method was used to create clean null mutants as described by Wang et al.(Wang *et al*, 2018) Specifically, a universal STOP-IN cassette with stop codons in all three reading frames was delivered by CRISPR-Cas9, using a single-strand DNA oligo containing the STOP-IN cassette and homology arms as the repair template. The STOP-IN cassette was inserted as close as possible to the start codon and was present in all the isoforms. Successful transgenic strains were identified by PCR gel electrophoresis and Sanger sequencing.

#### ROS in vivo imaging

*C. elegans* strain JV1, which expresses the H_2_O_2_ sensor HyPer, was used to measure ROS level *in vivo* in real-time. After the worms were trapped by *A. oligospora*, a 3 cm square LNM with a tested worm at the center was chunked, inverted, and mounted on the cover glass. Imaging was performed using an Andor Revolution WD system with a Nikon Ti-E automatic microscope and an Andor iXON Ultra 888 EMCCD camera at 1 fpm for 60 min. HyPer was excited by 488- and 405-nm lasers and Fiji analyzed HyPer ratiometric emissions at 525 nm and normalized the first frame’s 488/405 ratio to 1.

#### Antioxidant NAC treatment

To investigate the role of ROS in trapping-induced quiescence, we treated worms with the antioxidant N-acetylcysteine (NAC) before scoring pumping. The worms were treated in a 15 mL M9 solution containing NAC in a final concentration of 0.5mM in a 300 mL flask with continuous shaking at 150 rpm for 1 hour.(Gonzales-Moreno *et al*, 2023) NAC pre-treated worms were rinsed with M9 buffer before being added to the LNM plate for the experiment.

#### Data Availability

RNA sequencing data were deposited in the NCBI Gene Expression Omnibus (accession number: GSE243645) and are publicly available.

## Acknowledgments

We thank Dr. Henrik Bringmann, Dr. Chun-Ling Pan, and Dr. Chun-Hao Chen for sharing nematode strains. Some strains were provided by the CGC, which is funded by NIH Office of Research Infrastructure Programs (P40 OD010440). We thank WormBase. This work was supported by the Academia Sinica Investigator Award AS-IA-111-L02 to Y.-P.H. We would like to thank EMBO Global Investigator Network and Young Investigator Program for supporting Y.-P.H.

## Author contributions

Conceptualization, T.-H.L., H.-W.C., and Y.-P.H.; Formal Analysis, T.-H.L. and H.-W.C.; Investigation, T.-H.L. and H.-W.C.; Writing – Original Draft, T.-H.L., R.J.T., and Y.-P.H.; Writing – Review & Editing, T.-H.L., R.J.T., and Y.-P.H.; Visualization, T.-H.L.; Supervision, Y.-P.H.; Funding Acquisition, Y.-P.H.

## Declaration of interests

The authors declare no competing interests.

## Expanded View Figure legends

**Figure EV1.**
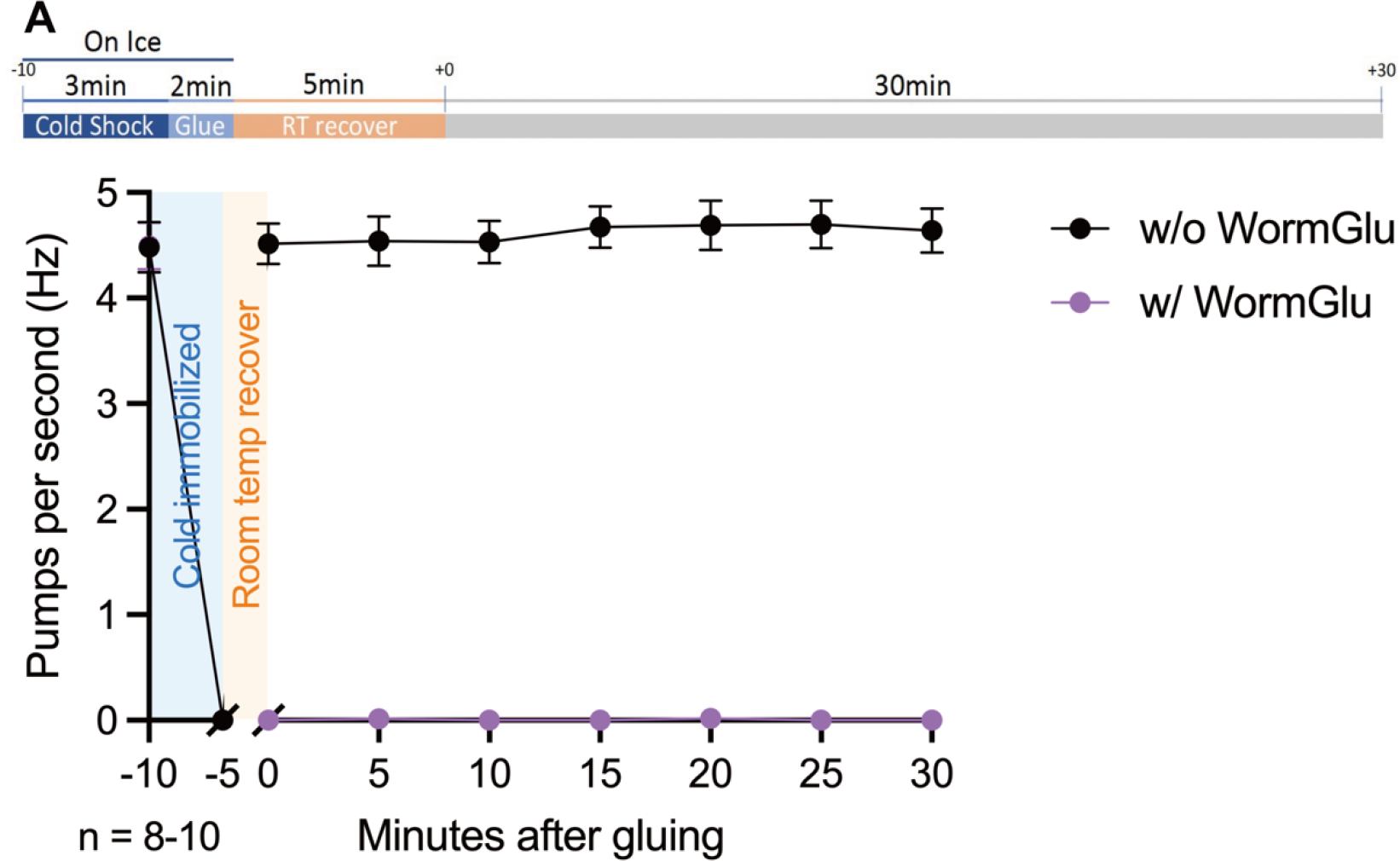
WormGlu adhesion-induced pumping quiescent (A) Schematic of the experimental procedure used to glue worms with WormGlu (upper) and traces showing pharyngeal pumping rates of worms with or without gluing (bottom).

**Figure EV2.**
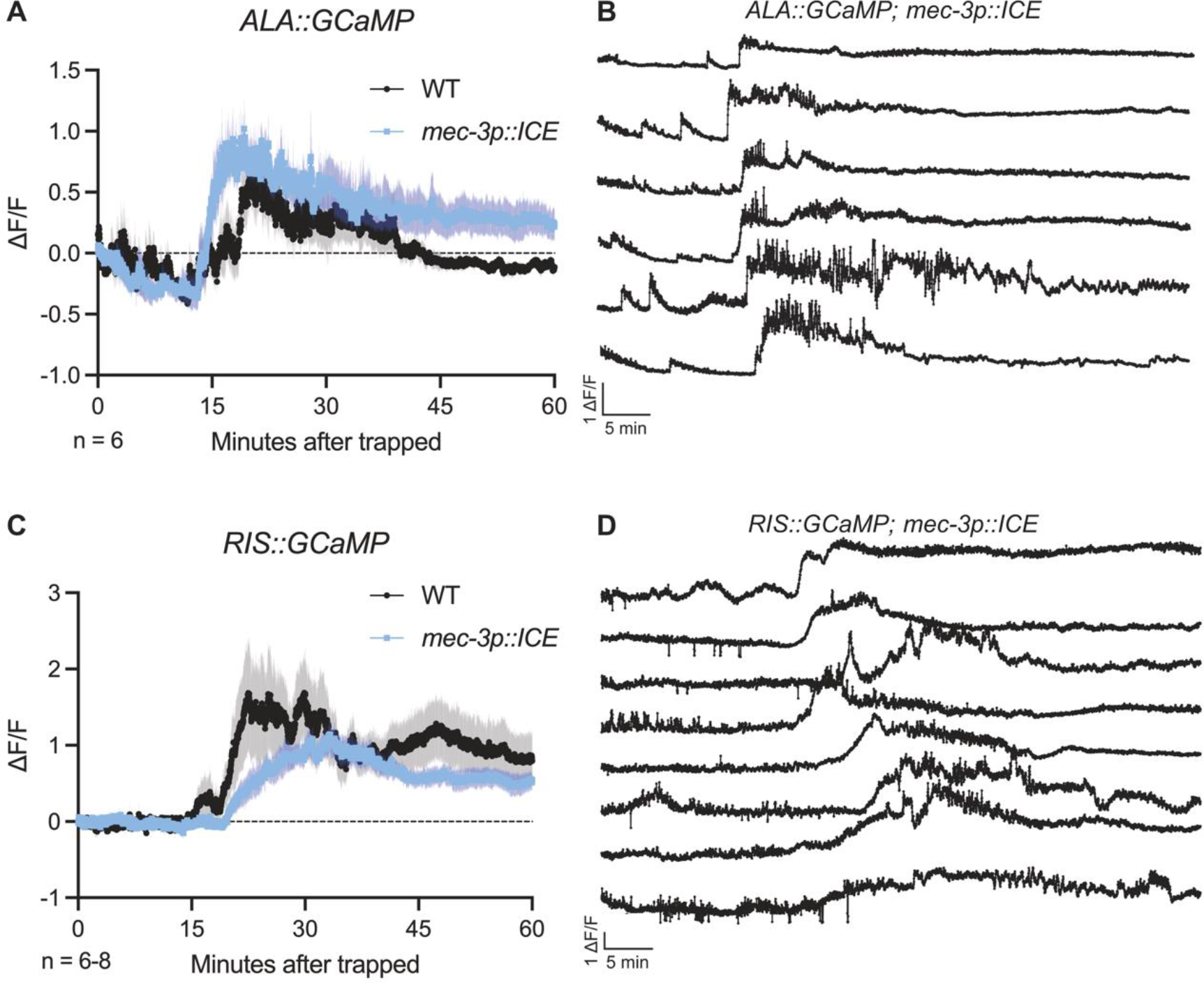
Mechanosensory pathways involved in sleep-promoting neuron activity patterns (A) Average trace of the ALA calcium response in genetically ablated mechanosensory mutants to *A. oligospora* trapping. WT data from Figure 2A. Mean ± SEM. (B) Individual traces of the ALA calcium response in genetically ablated mechanosensory mutants to *A. oligospora* trapping. (C) Average trace of the RIS calcium response in genetically ablated mechanosensory mutants to *A. oligospora* trapping. WT data from Figure 2C. Mean ± SEM. (D) Individual traces of the RIS calcium response in genetically ablated mechanosensory mutants to *A. oligospora* trapping.

**Figure EV3.**
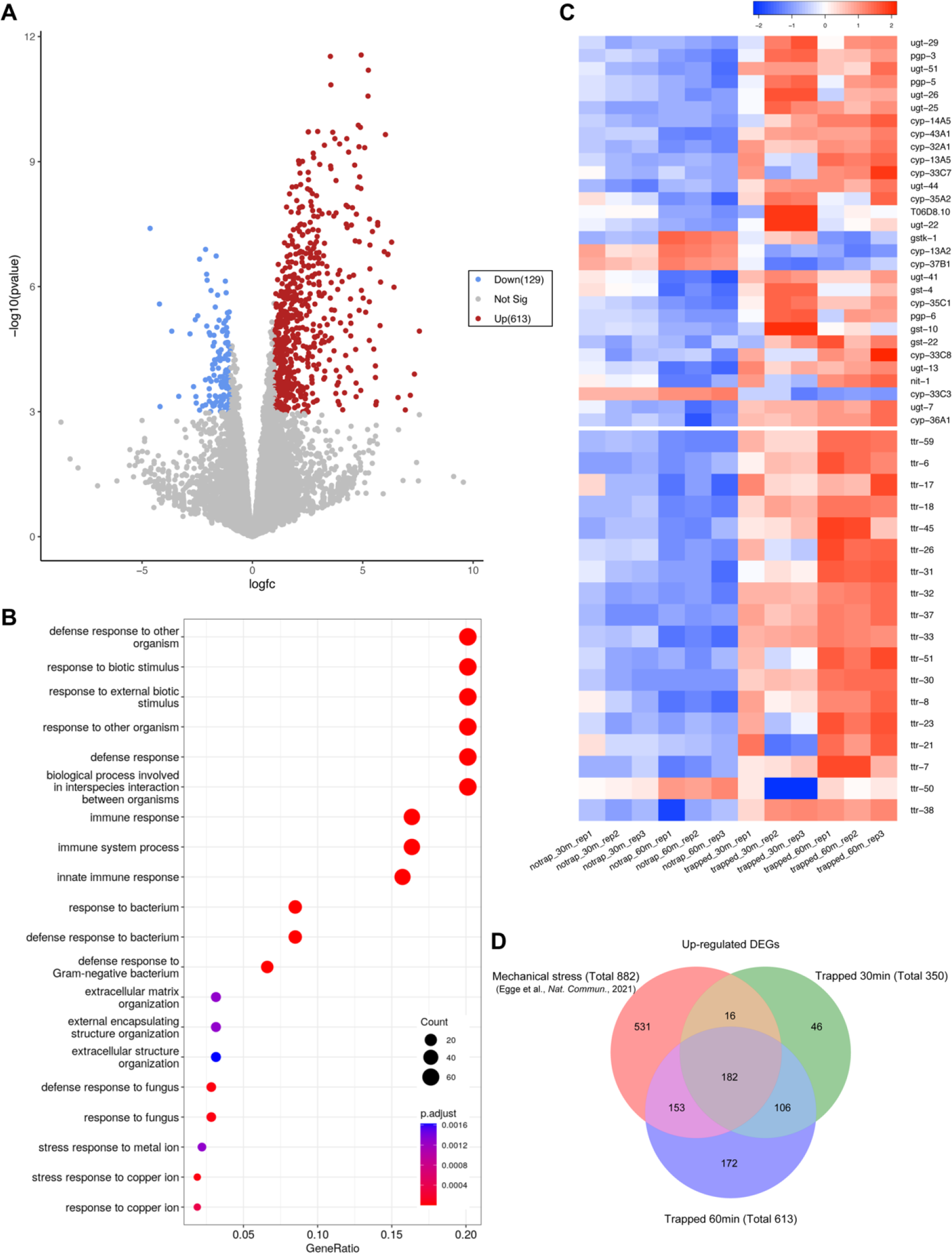
Transcriptomic analysis reveals up-regulation of XMEs and ttr genes (A) Volcano plot of RNA-seq analysis of 60 min trapped groups compared with the no-trap control. Threshold fold-change > 2; p-value < 0.05. (B) Plots of the gene enrichment analysis from GO molecular functions of 60 min trapped groups compared with the no-trap control. (C) Heat map showing the expression pattern of xenobiotic-related genes and *the ttr* gene family in different experimental groups. Normalized in Z-scale. (D) Venn diagram showing the relationship between upregulated DEGs in mechano-trauma-induced and *A. oligospora* trapping-induced RNA-seq data.

**Figure EV4.**
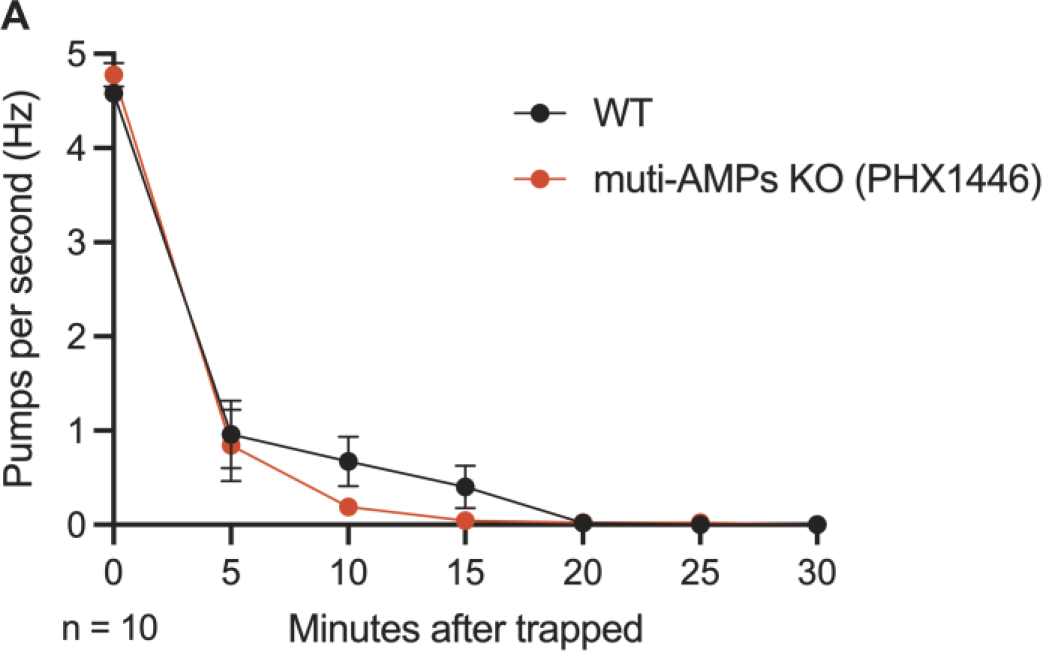
Antimicrobial peptide genes were not involved in trapping-induced pumping inhibition (A) Pharyngeal pumping rates of highly upregulated gene knockouts in response to *A. oligospora* trapping. PHX1446 carries knockouts of 19 members of the *nlp* and *cnc* peptide families. Mean ± SEM; Two-way ANOVA, Šidák’s multiple comparison test, comparing strain effects at each time point; all time points were non-significant.

## Expanded View Table and Movie

Table EV1. 182 core genes were consistently upregulated across all three RNA-seq datasets

**Movie EV1. *C. elegans* response to *A. oligospora* predation from struggling to quiescence**

This movie showed the *C. elegans* struggled in the first 17 minutes before entering a quiescent state. Red arrows indicate the trapped site. Time codes at the upper left indicate mm:ss. Speed up 240X. The scale bar represents 50µm.

